# Genome-Wide Identification and Characterization of *Fusarium circinatum*-Responsive lncRNAs in *Pinus radiata*

**DOI:** 10.1101/2021.07.21.453138

**Authors:** C. Zamora-Ballesteros, J. Martín-García, A. Suárez-Vega, J.J. Diez

## Abstract

One of the most promising strategies of Pine Pitch Canker (PPC) management is the use of reproductive plant material resistant to the disease. Understanding the complexity of plant transcriptome that underlies the defence to the causal agent *Fusarium circinatum*, would greatly facilitate the development of an accurate breeding program. Long non-coding RNAs (lncRNAs) are emerging as important transcriptional regulators under biotic stresses in plants. However, to date, characterization of lncRNAs in conifer trees has not been reported. In this study, transcriptomic identification of lncRNAs was carried out using strand-specific paired-end RNA sequencing, from *Pinus radiata* samples inoculated with *F. circinatum* at an early stage of infection. Overall, 13,312 lncRNAs were predicted through a bioinformatics approach, including long intergenic non-coding RNAs (92.3%), antisense lncRNAs (3.3%) and intronic lncRNAs (2.9%). Compared with protein-coding RNAs, pine lncRNAs are shorter, have lower expression, lower GC content and harbour fewer and shorter exons. A total of 164 differentially expressed (DE) lncRNAs were identified in response to *F. circinatum* infection in the inoculated versus mock-inoculated *P. radiata* seedlings. The predicted *cis*-regulated target genes of these pathogen-responsive lncRNAs were related to defence mechanisms such as kinase activity, phytohormone regulation, and cell wall reinforcement. Co-expression network analysis of DE lncRNAs, DE protein-coding RNAs and lncRNA target genes also indicated a potential network regulating pectinesterase activity and cell wall remodelling. This study presents the first analysis of conifer lncRNAs involved in the regulation of defence network and provides the basis for future functional characterizations of lncRNAs in relation to pine defence responses against *F. circinatum*.

## 1. Introduction

The major portion (98-99 %) of the transcribed genome comprises genetically inactive material known as non-coding RNA (ncRNA) (Lozada-Chávez *et al*. 2011). Among the ncRNA, the well-known housekeeping RNAs (transfer and ribosomal RNA) or small regulatory molecules including microRNAs (miRNAs), small nuclear RNAs (snRNAs) and small silencing RNAs (siRNAs) can be found (Bonnet *et al*. 2006). During the last decade, an heterogeneous class of ncRNA, long non-coding RNA (lncRNA), has emerged as another eukaryotic non-coding transcript class that had been largely ignored by molecular biologists (Tripathi *et al*. 2017). However, accumulating evidence supports that lncRNAs participate in many cellular processes by regulating gene expression in different manners (Quan *et al*. 2015). In this new and heterogeneous class, all transcripts greater than 200 nt in length that lack coding potential are included (Kapranov *et al*. 2007). Similar to protein-coding genes, lncRNAs are transcribed by RNA polymerase II, capped, polyadenylated and usually spliced (Quan *et al*. 2015). Some lncRNAs, termed *cis*-acting lncRNAs, regulate molecular processes around their transcription site, whereas *trans*-acting lncRNAs leave their transcription sites to exert their function elsewhere (Gil and Ulitsky 2020). LncRNAs are usually further sub-divided according to their function or based on their location and orientation with respect to the nearest protein-coding gene in the genome (Rai *et al*. 2019). Sense and anti-sense, intergenic as well as intronic (located into an intron) are the main groups for classifying the lncRNAs according to their genomic location (Ma *et al*. 2013). On the other hand, the reported functions for this class of transcripts differ substantially. Known mechanism of action including molecular signalling, decoys (binding to regulatory elements such as miRNAs blocking their molecular interaction), guides (directing specific RNA-protein complexes to specific targets) and scaffolds as central platforms for regulation, are associated to the majority of lncRNAs (Wang and Chang 2011).

The growing number of studies focusing on the interference of plant lncRNAs in different biological processes, including fertility, photomorphogenesis, wood formation, and biotic and abiotic stress, has demonstrated their important regulatory role in the transcription system (Chen *et al*. 2015; Liu *et al*. 2015; Sanchita *et al*. 2020). Some of these lncRNAs have been experimentally validated, most of them being from model plants. For example in *Arabidopsis*, two lncRNAs, COOLAIR and COLDAIR, have been shown to be crucial in the regulation of cold stress response (Swiezewski *et al*. 2009; Heo and Sung 2011). Likewise, DRIR lncRNA regulates the expression of a series of genes involved in drought and salt stress-responsive (Qin *et al*. 2017). The regulatory role of the lncRNA IPS1 has also been reported blocking the miRNA mir399 that suppress the expression of the gene responsible for the phosphate uptake (Franco-Zorrilla *et al*. 2007). Moreover, some lncRNAs associated with biotic stress have been characterized in plants. These included lncRNAs that regulate positively the expression of defence-related PR genes such as ELENA1, identified in *Arabidopsis* as a factor enhancing resistance against the pathogen *Pseudomonas syringae*, and lncRNA39026 that increases resistance against *Phytophthora infectans* in tomato (Seo *et al*. 2017; Hou *et al*. 2020). The biosynthesis or signalling of plant hormones have been altered by lncRNAs as well. In cotton plants, the silencing of two lncRNAs (GhlncNAT-ANX2 and GhlncNAT-RLP7) led to increased resistance to *Verticillium dahliae* and *Botrytis cinerea*, possibly due to the transcriptional induction of two lipoxygenases involved in the jasmonic acid defence signalling pathway (Zhang *et al*. 2018). In addition, overexpression of lncRNA ALEX1 in rice increased jasmonic acid levels enhancing resistance to the bacteria *Xanthomonas oryzae* pv. *oryzae* (Yu *et al*. 2020).

Next Generation Sequencing (NGS) technologies and computational methods have enabled a deeper study of the transcriptomic data and have been widely applied for the identification and characterization of plant lncRNAs (Tripathi *et al*. 2017). Recently, a number of lncRNAs involved in plant-pathogen interactions has been computationally predicted in non-model plants. In *Brassica napus*, 931 lncRNAs were identified in response to *Sclerotinia sclerotiorum* infection, one of them (TCONS_00000966) as antisense regulator of genes involved in plant defence (Joshi *et al*. 2016). Li *et al*. (2017) discovered *Musa acuminata* lncRNAs related to resistance against *Fusarium oxysporum* f. sp. *cubense* infection. Particularly, lncRNAs involved in the expression of pathogenesis-related proteins and peroxidases were mainly induced in the resistant cultivar, whereas lncRNAs related to auxin and salicylic acid signal transductions could predominantly be induced in the susceptible cultivar. In the Paulownia witches’ broom disease interaction, nine lncRNAs were predicted to target twelve genes based on a co-expression network model in the tree (Wang *et al*. 2017). In kiwifruit leaves infected by *P. syringae*, a weighted gene co-expression network analysis revealed a number of lncRNAs closely related to plant immune response and signal transduction (Wang *et al*. 2017). Likewise, Feng *et al*. (2021) identified 14,525 lncRNAs related to the walnut anthracnose resistance. This analysis showed that the target genes of the up-regulated lncRNAs were enriched in immune-related processes during the infection of the causal agent *Colletotrichum gloeosporioides*. These studies highlight the important role of lncRNAs in plant defence, thus further research is needed to decipher their function and interference in the transcriptomic system.

*Fusarium circinatum* is an invasive pathogen that causes the Pine Pitch Canker (PPC). This disease affects conifers, resulting in a serious economic and ecological impact on nurseries and pine stands (Wingfield *et al*. 2008). Since the first report in 1946 in North America, the presence of *F. circinatum* has been notified in 14 countries of America, Asia, Africa and Europe (Drenkhan *et al*. 2020). The long-distance dispersion as a result of globalization of plant trade and movement of contaminated soil and seed, represents the main pathway for new introductions of the pathogen into disease-free regions (Zamora-Ballesteros *et al*. 2019). The establishment of the disease in field is of great concern since no feasible measures are available to control or eradicate *F. circinatum* (Martín-García *et al*. 2019). Thus, the development of resistant genotypes through breeding and/or genetic engineering may be one of the most efficient PPC management strategy in the long-term (Gordon *et al*. 2015; Martín-García *et al*. 2019). In this context, several transcriptome analyses with the aim of unravelling molecular defence responses have provided detailed insights about the molecular mechanisms underlying disease progression in the *Pinus*-*F. circinatum* pathosystem. These studies have examined the response of hosts through a different degree of susceptibility, from highly susceptible (*Pinus radiata*, *Pinus patula*) to moderate (*Pinus pinaster*) and highly resistant (*Pinus tecunumanii*, *Pinus pinea*) (Visser *et al*. 2015, 2018, 2019; Carrasco *et al*. 2017; Hernandez-Escribano *et al*. 2020; Zamora-Ballesteros *et al*. 2021). However, the role of lncRNAs in the regulation of defence network in conifers has not been studied yet. In the present study, a strand-specific RNA-Seq has been conducted in order to characterize lncRNAs present in high susceptible *P. radiata* and elucidate how lncRNA expression profiles change in response to *F. circinatum* infection.

## 2. Material and methods

### 2.1. Inoculum preparation and inoculation trial

The *F. circinatum* isolate 072 obtained from an infected *P. radiata* tree in the North of Spain (Cantabria, Spain) was used. The isolate was cultured in Petri dishes containing PDA medium (Scharlab S.L., Spain) for a week at 25 °C. Then, to stimulate the sporulation of the fungus, four mycelial agar plugs were subcultured in an Erlenmeyer flask with 100 mL of PDB medium (Scharlab S.L., Spain) and incubated in an orbital shaker at 150 rpm during 48 hours at 25°C. Afterwards, the conidial suspension was adjusted with a haemocytometer at 10^6^ conidia mL^-1^ for the inoculation.

Six-month-old seedlings of *P. radiata* (Provenance: Galicia, Spain) with an approximate stem diameter of 2.5 ± 0.5 cm were inoculated on the stem by making a wound with a sterile scalpel and pipetting 10 µL of conidial suspension (Martin-Garcia *et al*. 2017). The same process was applied for the control seedlings that were mock-inoculated with sterilized distilled water. The inoculated wound was immediately sealed with Parafilm^®^ to prevent drying. Sixty seedlings were inoculated for each treatment (inoculation with pathogen and mock-inoculation). Plants were placed in a growth chamber at 21.5 °C with a 14-h photoperiod and kept for 67 days during which mortality rates were daily recorded.

The survival analysis based on the non-parametric estimator Kaplan-Meier (Kaplan and Meier 1958) was performed with the “Survival” package (Therneau 2020) to test the mortality of the plants. Survival curves were created with the “Survfit” function and the differences between the curves were tested with the “Survdiff” function. All analyses were performed using R software environment (R Core Team 2019).

### 2.2. RNA extraction and paired-end strand-specific sequencing

A piece of the stem from the upper part of the inoculation point (*ca*. 1 cm length) was sampled at four days post-inoculation (dpi) for the transcriptomic analysis. The harvested tissues were immediately frozen in liquid nitrogen and ground to a fine powder using a mortar and pestle. RNA extraction were performed using the Spectrum™ Plant Total RNA Kit (Sigma Aldrich, USA) following the manufacturer’s protocols including the optional on-column DNase 1 digestion (DNASE10-1SET, Sigma-Aldrich, St. Louis, MO, USA). After RNA extraction, samples were transferred to RNase- and DNase-free tubes (Axygen^®^, USA) and stored at −80 °C. The concentration and purity of the RNA extracted were measured using the Multiskan GO Spectrophotometer (A_260_/A_280_ ≥ 1.8, A_260_/A_230_ ≥ 1.8 and concentration > 50 ng/µl; Thermo Fisher Scientific, Waltham, MA, USA). RNA integrity was checked by agarose gel electrophoresis (1% TAE).

Six biological replicates of inoculated and three of mock-inoculated treatment were sent to Macrogen Co. (Seoul, South Korea) for sequencing. Sequenced samples showed a RNA integrity number (RIN) ≥ 7 measured by an Agilent 2100 Bioanalyzer. The strand-specific RNA-Seq libraries were constructed using the Illumina TruSeq Stranded mRNA protocol with polyadenylated mRNAs and lncRNAs enrichment and an insert size of 300 bp (150×2 paired-end reads). Sequencing was performed on the Illumina NovaSeq 6000 Sequencing System (Illumina Inc., USA).

### 2.3. Genome mapping and reference-based transcriptome assembly

All sequenced libraries were assessed for quality control using FastQC v.0.11.9 (Andrews 2012) and trimmed for Illumina adaptor sequences and low-quality base-calls using Trimmomatic v.0.38 (Bolger *et al*. 2014). The trimmed reads with high quality were then aligned to the *Pinus taeda* reference genome sequence (Pita_v2.01; Treegenes database, Wegrzyn *et al*. 2008) using HISAT2 v.2.0.0 (Kim *et al*. 2015) with parameters “--known-splicesite-infile”, “--dta” and “--rna-strandness RF”. In order to ensure the presence of *F. circinatum* biomass in the samples, the reads were also mapped to its publically available genome sequence (accession number JAGGEA000000000). The SAM files from the pine mapping were processed with the SAMtools utility (Li *et al*. 2009) for converting to binary alignment map (BAM) format, sorting by coordinates and removing duplicates. The transcripts for each sample were reconstructed separately by StringTie v.2.1.4 (Pertea *et al*. 2015) using the “-G option” with the annotation file of *P. taeda* (Pita.2_01.entap_annotations.tsv; Treegenes database, Wegrzyn *et al*. 2008). This file was previously fixed with Gffread utility v.0.12.1 (Pertea and Pertea 2020) for the correct understanding by StringTie program. After the transcriptome assembly, the nine resulting GTF files were merged to generate a non-redundant set of transcripts with unique identifiers using the StringTie “-merge” parameter, where only transcripts with expression levels > 0.1 fragment per kilobase of exon per million mapped reads (FPKM) were included. Finally, this newly experiment-level transcriptome was further compared with the *P. taeda* reference annotation GTF file (Pita_v2.01; Treegenes database, Wegrzyn *et al*. 2008) using the software Gffcompare v.0.12.1 (Pertea and Pertea 2020), classifying transcripts in different class codes according to their nature/origin.

### 2.4. LncRNAs identification

Based on all the assembled transcripts, the known transcripts marked with the class code “=” were excluded before conducting the potential long non-coding RNAs identification. The remaining transcripts were subjected to the coding potential predictor FEELnc v.0.2 tool (Wucher *et al*. 2017) as well as several filters to ensure reliability of lncRNAs. Firstly, the FEELnc filter module was used to remove short transcripts (< 200 nt) and retain monoexonic transcripts with antisense localization. After that, the sequences of the resulting transcripts were extracted with Gffread v.0.12.1 (Pertea and Pertea 2020) and the fasta file output was piped to the Eukaryotic Non-Model Transcriptome Annotation Pipeline (EnTAP) v.0.9.2 (Hart *et al*. 2020) for transcript annotation. Briefly, GeneMarkS-T v.5.1 (Tang *et al*. 2015) was used for open reading frame (ORF) prediction and the sequence aligner DIAMOND v.1.9.2 (Buchfink *et al*. 2015) conducted the similarity search with default settings (E-value < 10^-5^) using the NCBI non-redundant protein database (release-201). After that, the assignment of protein domains (Pfam), Gene Ontology (GO) terms and KEGG pathways was performed using EggNOG v.1.0.3 (Huerta-Cepas *et al*. 2016). Finally, EnTAP filtered contaminants to retain only high-quality transcripts. Subsequently, the FEELnc codpot module was used with the shuffling mode to calculate a coding potential score (CPS) for the un-annotated transcripts using a random forest algorithm trained with multi k-mer frequencies and relaxed ORFs. The specificity threshold was set at 0.95 in order to increase the robustness of the final set of novel lncRNAs. The remaining transcripts were designated as lncRNAs and further classified according to the ‘Gffcompare’ output as long intergenic non-coding RNAs (lincRNAs) categorized with class code ‘u’, long non-coding natural antisense transcripts (lncNAT) from the class code ‘x’, and intronic transcripts that were those with class code ‘i’ (Budak *et al*. 2020).

In order to investigate the conservation of the pine lncRNAs, two recently released and updated databases of known plant lncRNAs were used (Rai *et al*. 2019). All the transcripts designated as lncRNA were aligned against CANTATA database (Szcześniak *et al*. 2019) and GreeNc database (Gallart *et al*. 2016) using the blastn algorithm (E-value <10^-5^) of the BLAST v.2.9.0 software suite (Kozomara *et al*. 2019; Kalvari *et al*. 2021). Moreover, the transcripts were also aligned to the Rfam (version 14.1) and miRBase (version 21) non-coding RNA databases with designated threshold value (E-value <10^-5^) using the blastn algorithm in order to detect housekeeping non-coding RNAs including transfer RNA (tRNAs), ribosomal RNA (rRNAs) and snoRNAs, and miRNA precursors.

### 2.5. Differential expression analysis

StringTie together with the “-e” parameter was employed to estimate expression for all transcripts of the experiment-level transcriptome (Pertea *et al*. 2015). The output file was reformatted using the “prepDE.py” script for further expression analysis (CCB 2019). DESeq2 v.1.24.1 (Love *et al*. 2014) was used to identify differentially expressed (DE) lncRNA transcripts based on the matrix of the estimated counts. Differentially expressed genes (DEG) were identified equally. The pairwise comparison of inoculated and control plants were evaluated using Wald tests. To visualize the similarity of the replicates and identify any sample outliers, the principal component analyses (PCA) was constructed using the rlog-transformed expression values. Transcripts were considered as differentially expressed if the adjusted p-values (padj) for multiple testing, using Benjamini–Hochberg to estimate the false discovery rate (FDR) (Benjamini and Hochberg 1995), was less than 0.05 and the |log2 (Fold Change)| ≥ 1.

### 2.6. Potential target gene prediction and functional enrichment

Based on the genome location of the lncRNAs relative to the neighbouring genes, the nearest protein-coding genes transcribed within a 10 kb window upstream or 100 kb downstream were considered as potential *cis*-regulated target genes. These genes were identified using the FEELnc classifier module (Wucher *et al*. 2017) and annotated using the EnTAP pipeline (Hart *et al*. 2020) as described above but implemented with the RefSeq complete protein database (release-201) and the UniProtKB/Swissprot database (release-2020_05).

Functional enrichment analysis of the target genes associated with the DE lncRNAs was conducted. DE lncRNA transcripts were divided into up- and down-regulated subsets for efficient functional analysis (Hong *et al*. 2014). Using all genes as background, GO and KEGG enrichment analysis were conducted by GOSeq v.1.38.0 based on the Wallenius non-central hyper-geometric distribution that allows the adjustment for transcript length bias (Young *et al*. 2010). The GO terms and KEGG pathways with corrected p-values lower than 0.05 were considered to be enriched in the group. Redundant gene ontology categories were parsed using Revigo (Supek *et al*. 2011).

### 2.7. Co-expression analysis and identification of hub genes

In order to predict the co-expression modules and determine the GO terms that differentiate the transcriptome induced by *F. circinatum*, a weighted gene co-expression network analysis approach implemented in the R-based Co-Expression Modules identification Tool (CEMiTool) package v.1.8.3 (Russo *et al*. 2018) was conducted in R software. Network analysis was carried out on the expression data for three gene sets: DE lncRNAs, DEGs and targeted genes predicted by FEELnc. A variance stabilizing transformation (vst) was used and transcripts were filtered to reduce correlation between variance and gene expression. The Spearman’s method was used for calculating the correlation coefficients and a soft thresholding power (β) of 6 was selected. The co-expressed modules were subjected to over-representation analysis (ORA) based on the hypergeometric test (Yu *et al*. 2012) using the GO terms to determine the most significant module functions (q-value ≤ 0.05) (Russo *et al*. 2018). Moreover, genes with the highest connectivity, known as hub genes and considered functionally-important genes (Tahmasebi *et al*. 2019) were identified in each module.

## 3. Results

### 3.1. Disease monitoring

The survival analysis revealed clear significant differences between the inoculation and control conditions (χ^2^ = 116, p < 0.001). At 10 dpi all seedlings inoculated with *F. circinatum* showed symptoms of PPC (resin and/or necrosis at the stem and wilting) and started to die at 33 dpi (**Figure 1**). By the end of the experiment, 92.2% of the inoculated seedlings had died. No mortality was recorded for control seedlings.

**Figure 1.**
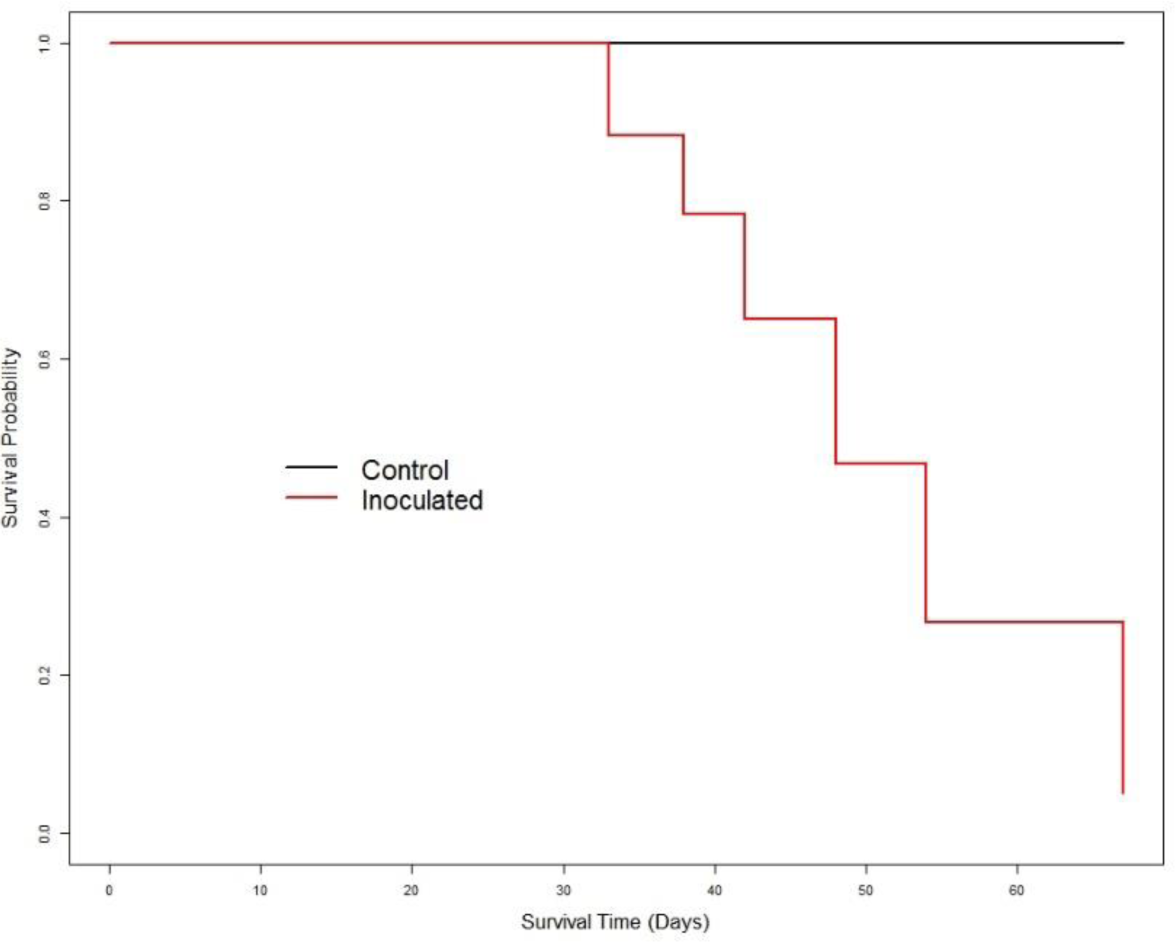
Survival probability plot for *P. radiata* seedlings inoculated with *F. circinatum* and mock-inoculated, determined using the Kaplan-Meier estimate of the survival function.

### 3.2. Deep sequencing and transcripts assembly

High-throughput strand-specific RNA-Seq of nine libraries constructed from stem tissue of *P. radiata* inoculated with *F. circinatum* and mock-inoculated were analysed. Raw data of the experiment have been deposited at the NCBI under the SRA numbers SRR15100123-31 (BioProject PRJNA742852). Almost 590 million 150-base pair-end reads on polyadenylated selected (polyA) RNAs were generated by the Illumina platform. RNA-Seq reached average depths of *ca*. 65.5 million reads (55 to 84 million reads) (**Table S1**). After adapter and low-quality nucleotides trimming, an average of 78% of paired reads and 11% of mates from broken pairs were retained. Approximately 74.21% and 70.33% of reads from inoculated and mock-inoculated libraries successfully mapped to the *P. taeda* reference genome, respectively (**Table S1**). Considering the infected samples, the average of 2.63% reads mapped to the *F. circinatum* genome confirmed the presence of the pathogen.

**Table 1.** Number of unknown transcripts of *P. radiata* associated to a class code according to GffCompare software classification.

| Class code | After assembly |  | LncRNAs predicted |  | Description<sup>1</sup> |
| --- | --- | --- | --- | --- | --- |
|  | Transcript no. | % | Transcript no. | % |  |
| **x** | 902 | 0.71 | 446 | 3.35 | Overlapping an exon of an annotated gene at the opposite strand |
| **i** | 1,178 | 0.92 | 383 | 2.88 | Fully contained in a known intron |
| **y** | 500 | 0.39 | 189 | 1.42 | Contains a reference gene within its intron |
| **p** | 516 | 0.4 | 0 | 0 | Adjacent to the 5' end of an annotated gene at the same strand |
| **u** | 45,705 | 35.8 | 12,280 | 92.32 | Intergenic region |

<sup>1</sup>Brief explanation of the class codes.

Nine high-depth transcriptomes were generated. Six of them were reconstructed from *P. radiata* inoculated with *F. circinatum*, and the other three were generated from the mock-inoculated seedlings. After merging all of them, the unique transcriptome assembled were composed of 87,427 loci and 127,677 transcripts, with 43.1% GC content. A total of 51,212 (40.11%) transcripts were shared with the reference annotation file (Pita_v2.01.gtf) and discarded for lncRNA detection analysis since these transcripts were known as protein-coding RNAs. The remaining 76,465 transcripts were further categorized into different class codes according to its relationship with its closest reference transcript (**Table 1**).

### 3.3. Genome-wide identification and characterization of pine lncRNAs

The 76,465 total unknown transcripts were subjected to several sequential filter steps to obtain the lncRNA transcripts (**Figure 2**). A total of 13,312 lncRNAs (length ≥ 200 nt, ORF coverage < 50%, and potential coding score < 0.5) and 47,473 potential new isoforms were obtained at the end of the pipeline. Using the FEELclassifier module, the class distributions of the pine lncRNAs was performed according to their location relative to the nearest protein-coding gene based on the reconstructed transcriptome. The majority of the lncRNAs were lincRNAs with 12,291 (92.3%) transcripts, followed by lncNAT with 445 (3.3%) transcripts and 383 (2.9%) intronic transcripts. In addition, 25 lncRNAs were also identified as known miRNA precursors belonging to 10 miRNA families being the most represented MIR160, MIR159 and MIR1314. The Rfam and miRBase analyses also allowed the identification of 174 transcripts that were found to be distributed among 32 conserved RNA families including rRNA, tRNA, histones and several snoRNAs (**Table S2-S3**).

**Figure 2.**
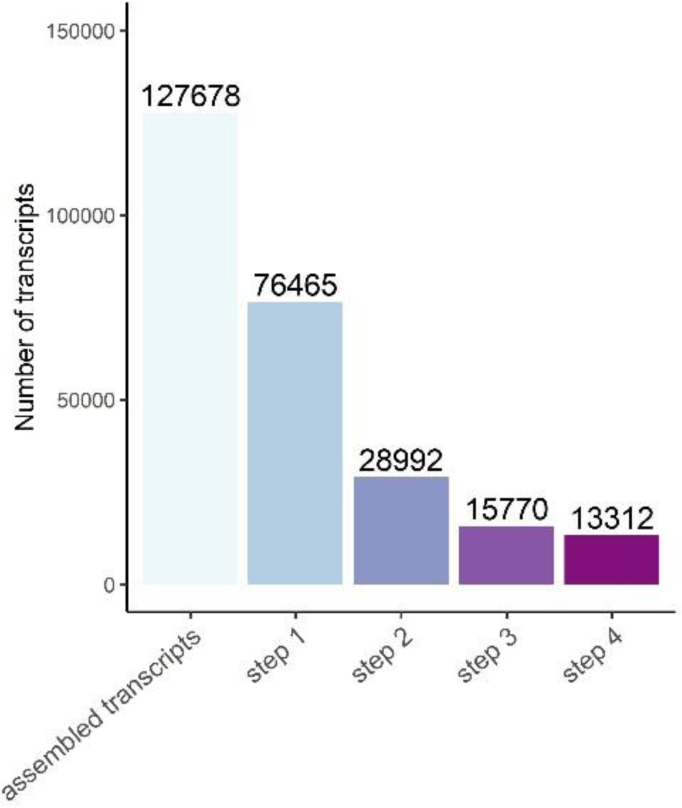
Statistics of candidate lncRNA transcripts. Step 1: known protein-coding transcripts were filtered out. Step 2: transcripts with length ≥ 200 bp and with at least two exons (including monoexonic transcripts with antisense localization) were selected. Step 3: transcripts annotated with EnTAP program were filtered out. Step 4: with a coding potential score lower than < 0.5 were retained.

**Table 2.** Candidate target genes predicted to interact with DE lncRNA transcripts.

| LncRNA | Log<sub>2</sub>FC | Targeted gene | Log<sub>2</sub>FC | Direction | Type | Distance | Subtype | Location | Description of targeted gene |
| --- | --- | --- | --- | --- | --- | --- | --- | --- | --- |
| lncRNAPiRa.44237.18 | 9.76 | PITA_00496 |  | antisense | intergenic | 41417 | convergent | downstream | Pentatricopeptide repeat-containing protein At4g13650 |
| lncRNAPiRa.64325.1 | 3.15 | PITA_01014 |  | antisense | genic | 0 | containing | exonic | Transcript with domain: DUF4228 |
| lncRNAPiRa.32343.2 | 11.3 | PITA_13284 |  | antisense | intergenic | 51151 | convergent | downstream | CYCD2 |
| lncRNAPiRa.35491.1 | 9.18 | PITA_33574 |  | antisense | intergenic | 3155 | convergent | downstream | Leaf rust 10 disease-resistance locus receptor-like protein kinase-like 1.2 isoform X1 |
| lncRNAPiRa.42942.2 | 8.54 | PITA_15284 |  | antisense | intergenic | 92774 | divergent | upstream | Ribosomal RNA methyltransferase FtsJ domain-containing protein |
| lncRNAPiRa.22160.1 | 9.5 | PITA_12411 | 6.58 | sense | intergenic | 9711 | same_strand | downstream | Pectin methylesterase 17 |
| lncRNAPiRa.31525.1 | 9.63 | PITA_31792 |  | antisense | intergenic | 35358 | convergent | downstream | Transcript with domain: Myb_DNA-binding |
| lncRNAPiRa.79902.12 |  | PITA_05666 | 10.1 | sense | genic | 0 | containing | intronic | PDC1 |
|  | 7.12 | PITA_12210 | 11.9 | sense | intergenic | 87 | same_strand | downstream | PDC1 |
| lncRNAPiRa.70333.4 | 7.9 | PITA_01539 |  | sense | genic | 0 | nested | intronic | Uclacyanin 1 |
| lncRNAPiRa.51697.3 | 3.35 | PITA_34628 |  | sense | genic | 0 | containing | intronic | Transcript with domain: Peptidase_S28, Peptidase_S9 |
| lncRNAPiRa.61651.3 | 8.53 | PITA_42898 |  | antisense | intergenic | 7280 | convergent | downstream | PAR1 |
| lncRNAPiRa.33277.3 | 3.35 | PITA_08467 | 3.54 | sense | intergenic | 376 | same_strand | upstream | PAR1 |
| lncRNAPiRa.45077.2 | 8.14 | PITA_26106 |  | antisense | intergenic | 87766 | convergent | downstream | Purple acid phosphatase |
| lncRNAPiRa.23041.2 | 9.27 | PITA_28262 |  | sense | intergenic | 9586 | same_strand | downstream | Pectin methylesterase 17 |
| lncRNAPiRa.85490.1 | 6.99 | PITA_28228 |  | antisense | intergenic | 85760 | divergent | upstream | unknown [Picea sitchensis] |
| lncRNAPiRa.47042.1 | 7.53 | PITA_13092 |  | sense | intergenic | 31121 | same_strand | upstream | Transcript with domain: PP2C |
| lncRNAPiRa.19024.1 | 5.58 | PITA_42377 | 5.17 | sense | intergenic | 542 | same_strand | downstream | Non-symbiotic hemoglobin 1 (HB) |
| lncRNAPiRa.25700.7 |  | PITA_23327 |  | sense | genic | 0 | containing | intronic | Peptidase S9 |
|  | 3.79 | PITA_25465 |  | sense | intergenic | 541 | same_strand | downstream | Prolyl endopeptidase |
| lncRNAPiRa.25968.1 | 2.91 | PITA_42840 | 4.85 | sense | intergenic | 33267 | same_strand | downstream | Transcript with domain: USP |
| lncRNAPiRa.29628.1 | 6.8 | PITA_10474 |  | sense | intergenic | 812 | same_strand | downstream | Transcript with domain: Glycolytic-Fructose-bisphosphate aldolase class-I |
| lncRNAPiRa.80857.1 | 6.78 | PITA_28959 |  | antisense | genic | 0 | nested | intronic | ALN |
| lncRNAPiRa.29753.1 | 7.2 | PITA_38537 | 6.4 | sense | intergenic | 69405 | same_strand | downstream | leaf rust 10 disease-resistance locus receptor-like protein kinase-like protein 2.4 |
| lncRNAPiRa.33098.2 | 6.8 | PITA_43179 | 5.01 | sense | intergenic | 6072 | same_strand | downstream | 4-coumarate-CoA ligase, partial (4CL3) |
| lncRNAPiRa.80336.1 | 4.89 | PITA_17252 |  | antisense | intergenic | 47603 | convergent | downstream | Protein chromatin remodeling 24 |
| lncRNAPiRa.61651.4 | 5.82 | PITA_42898 |  | antisense | intergenic | 7280 | convergent | downstream | unknown [Picea sitchensis] |
| lncRNAPiRa.64704.5 | 6.91 | PITA_22879 |  | sense | intergenic | 6023 | same_strand | downstream | Lambda class glutathione S-transferase (GSTL1) |
| lncRNAPiRa.75647.1 | 5.93 | PITA_04032 |  | antisense | intergenic | 8949 | divergent | upstream | Transcript with domain: RRM_1 |
| lncRNAPiRa.85000.6 | 9.91 | PITA_16990 |  | sense | intergenic | 5468 | same_strand | upstream | Ethylene receptor 2 (ETR2) |
| lncRNAPiRa.33190.1 | 7.85 | PITA_44567 |  | sense | intergenic | 66345 | same_strand | downstream | Transcript with domain: EamA |
| lncRNAPiRa.31184.1 | 2.25 | PITA_16807 |  | antisense | intergenic | 55755 | divergent | upstream | Transcript with domain: LEA_3 |
| lncRNAPiRa.78332.11 | 2.65 | PITA_41139 | 4.04 | sense | genic | 0 | overlapping | intronic | CBS domain-containing protein cbscbpsb3 |
| lncRNAPiRa.42813.1 | 9.29 | PITA_02986 |  | sense | intergenic | 190 | same_strand | downstream | Hypothetical protein 0_9919_01, partial [Pinus taeda] |
| lncRNAPiRa.78487.3 | 9.39 | PITA_28133 |  | antisense | intergenic | 33344 | convergent | downstream | Transcript with domain: zf-CCCH |
| lncRNAPiRa.84511.1 | 4.53 | PITA_13110 | 6.3 | sense | intergenic | 82 | same_strand | downstream | Transcript with domain: Cellulase |
| lncRNAPiRa.62823.1 | -9.14 | PITA_01229 |  | sense | intergenic | 76300 | same_strand | downstream | UBA52 |
| lncRNAPiRa.83146.2 | -7.43 | PITA_05626 |  | sense | intergenic | 69363 | same_strand | downstream | Pyridoxal kinase-like protein isoform X1 |
| lncRNAPiRa.38350.3 | -6.97 | PITA_18454 |  | sense | intergenic | 494 | same_strand | downstream | CC-NBS-LRR resistance-like protein |

The average length of protein-coding transcripts (1,200 bp) was higher than that of lincRNAs (750 bp), lncNATs (452 bp) and intronic lncRNAs (565 bp). However, while most of lncNATs and intronic lncRNAs showed short lengths (300 bp), lincRNAs and protein-coding transcripts exhibited a similar trend of length distribution (**Figure 3A**). Overall, the size distribution of the lncRNAs ranged from 200 to 7,393 bp, with the majority of these transcripts ranging from 200 to 400 bp. Differences in the analysis of the exon number were also found. While the lncRNAs showed an average exon number of 2.5, the protein-coding transcripts had 4.1 exons (**Figure 3B**). This analysis also revealed that two-exon transcripts were the most represented in this study. The highest ratio of two-exon transcripts was found in lncNATs (77.3%) and intronic lncRNAs (75.7%), followed by lincRNAs (66.9%). In the group of protein-coding transcripts, the ratio of two-exon transcripts was not so high (32%). Regarding the exon length, similarly to the transcript length, the exons belonging to the lncNAT and intronic lncRNA transcripts showed shorter lengths (100-300 bp) than those belonging to protein-coding transcripts (**Figure 3C**). Once again, the distribution of the exon lengths from the lincRNA transcripts was similar to that of protein-coding transcripts.

**Figure 3.**
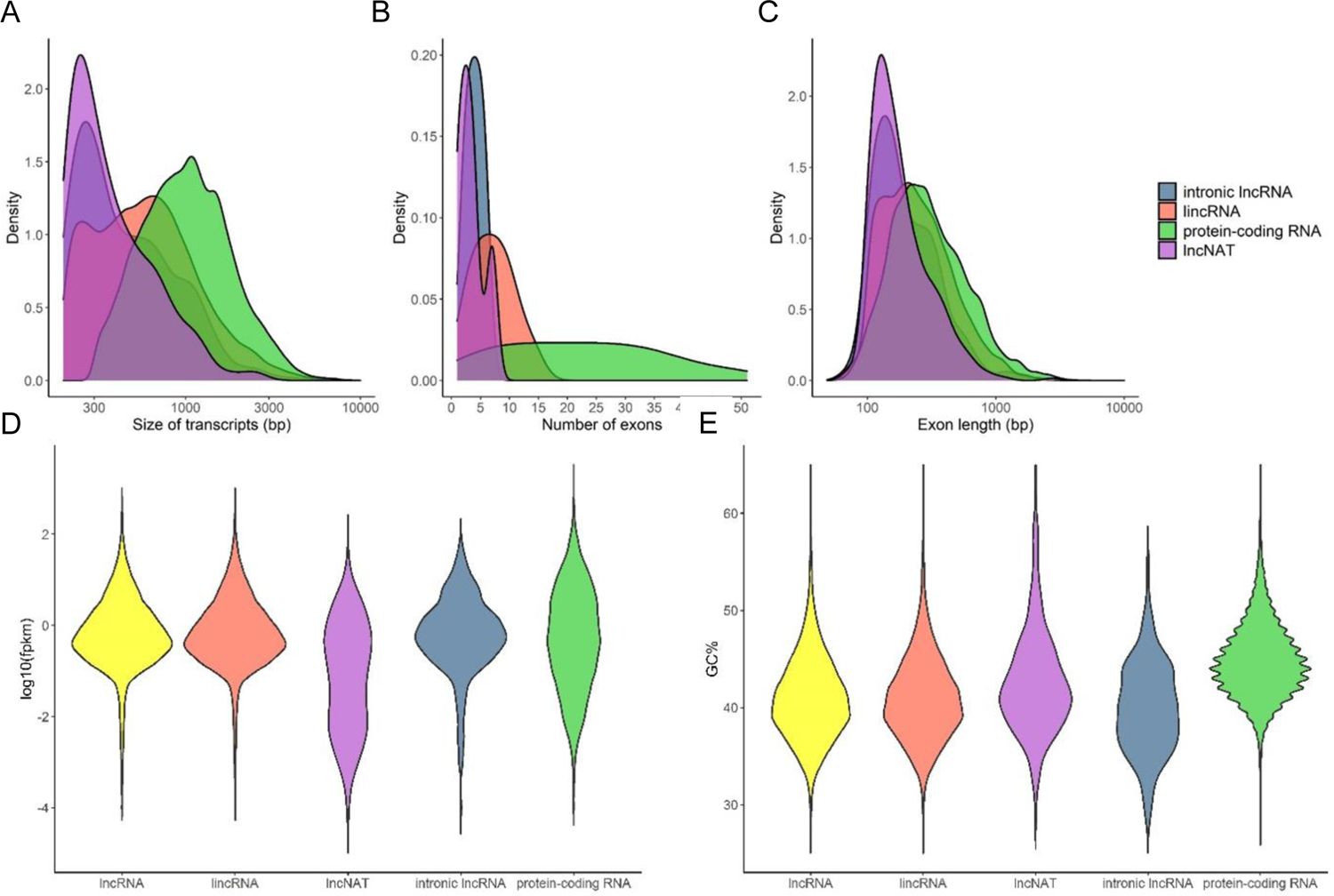
The characterization lncRNA transcripts showed differences with the characteristics of protein-coding transcripts in *P. radiata*. (A) Transcript size distribution for lincRNAs, lncNATs, intronic lncRNAs and protein-coding RNAs. (B) Number of exons per transcript for lincRNAs, lncNATs, intronic lncRNAs and protein-coding RNAs. (C) Exon size distributions for lincRNAs, lncNATs, intronic lncRNAs and protein-coding RNAs. (D) FPKM distribution of lncRNAs and protein-coding RNAs. (E) GC content of lncRNAs and protein-coding RNAs.

The average expression levels of lncRNAs in terms of FPKM was lower (3.3) than those of protein-coding transcripts (5.6) (**Figure 3D**). In addition, the GC content in lncRNAs (41%) was slightly lower than that in protein-coding transcripts (44.8%), showing the intronic lncRNA transcripts the lowest percentage (**Figure 3E**).

All the lncRNA transcripts were aligned against the known lncRNAs of 10 different plant species from the CANTATA database: *Chenopodium quinoa*, *Brassica napus*, *Malus domestica*, *Zea mays*, *Arabidopsis thaliana*, *Oryza rufipogon*, *Vitis vinifera*, *Populus trichocarpa*, *Prunus persica* and *Ananas comosus*. Likewise, known lncRNAs of all plant species present in the GreeNc database, except those species already examined with the CANTATA database, were confronted with the lncRNAs of *P. radiata*. A number of 1,131 (8.6%) lncRNAs were conserved across the ten species of CANTATA (**Table S4**). In addition, a total of 1,421 (10.8%) lncRNA transcripts, corresponding to known lncRNA genes from the GreeNc database (**Table S5**), were obtained. Therefore, 2,552 (19.3%) lncRNAs showed homology with known lncRNAs from other plant species. The highest homology ratio (number of hits of pine lncRNAs with those of each plant species to the total number of lncRNAs of each plant species) was observed with the woody plant *P. trichocarpa* (5.03%) (**Figure S1**).

### 3.4. Differentially expressed analysis in response to *F. circinatum* infection and prediction of candidate target genes

The expression changes of lncRNAs between the *P. radiata* seedlings inoculated with *F. circinatum* and controls were analysed. The PCA allowed to identify two sample outliers among the pathogen-inoculated condition that were discarded for the differential expression analysis (**Figure S2**). A total of 164 lncRNA transcripts were identified as differentially expressed (p-value < 0.05, log_2_ (|Fold-change|) ≥ 1) under the pathogen infection, 146 of which were up-regulated and 18 down-regulated (**Table S6-S7**). Among the DE lncRNAs, 157 transcripts were lincRNA and the remainder were two intronic lncRNAs, one lncNATs, and four lncRNAs containing a coding-protein in its intron. DE lncRNAs were clustered in a heat map in order to visualize the expression pattern of both conditions of the analysis (**Figure 4**). On the other hand, 2,369 protein-coding RNA were up-regulated and 189 down-regulated by the pathogen infection (**Table S8-S9**).

**Figure 4.**
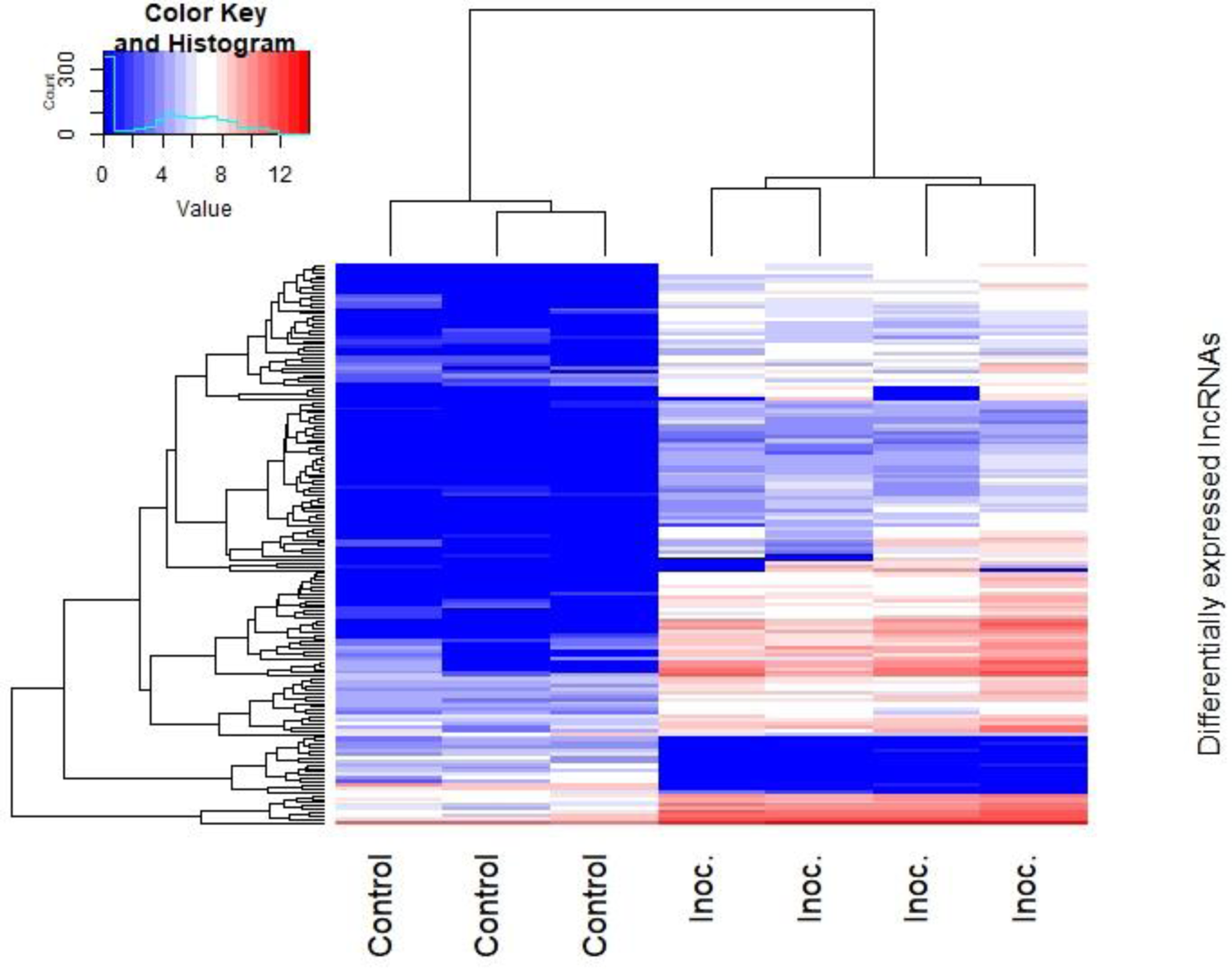
The hierarchical clustering plot shows the scaled expression levels of the differentially expressed lncRNAs of *P. radiata* in response to *F. circinatum*. Different columns represent different libraries, and different rows represent the differentially expressed lncRNAs. Red: relatively high expression; Blue: relatively low expression.

### 3.5. Analysis of lncRNAs *cis*-interacting genes

To predict the role of *cis*-acting lncRNAs of *P. radiata* in response to *F. circinatum*, the protein-coding transcripts located within a 10 kb window upstream and 100 kb downstream were investigated. A total of 4,268 lncRNA–mRNA interaction pairs were recorded by the FEELnc classifier module (**Table S10**). However, one lncRNA could have more than one target gene, and a target gene could be the target of one or more lncRNAs. In fact, a number of 2,760 candidate *cis* target genes were observed for 3,750 lncRNAs, of which 3,342 had a single candidate target gene and 408 lncRNAs had multiple interactions. The maximum number of target genes for a single lncRNA was five, which was reached by seven lncRNAs (**Table S11**). Moreover, the 73% of the 2,760 candidate target genes were targeted by one lncRNA, while one candidate target gene could be targeted by up to 30 different lncRNAs.

In total, 39 candidate target genes were predicted for the 37 DE lncRNAs (**Table 2**). The function prediction of these DE lncRNAs was based on the functional annotation of their nearby target genes. Among these targeted genes, there were genes encoding for receptor-like protein kinases (RLKs), enzymes associated to the cell-wall reinforcement and lignification (pectin methylesterases inhibitor, uclacyanin and 4-coumarate-CoA ligase), and enzymes involved in the attenuation of oxidative stress (glutathione S-transferase). One RLK that was predicted to be targeted by the up-regulated lncRNAPiRa.29753.1 was, in turn, induced by the pathogen infection. Two pectin methylesterases (PME) were predicted to be regulated by lncRNAPiRa.23041.2 and lncRNAPiRa.22160.1 transcribed in the same orientation in a downstream location. One of the targeted PME was DE by the pathogen infection, whereas the other PME did not. Moreover, the coding region for 4-coumarate-CoA ligase 3 (4CL3) targeted by lncRNAPiRa.33098.2 was also present among the DEGs of the coding RNAs analysis. One gene harbouring the DNA-binding motif MYB, a transcription factor with a role in plant stress tolerance, was potentially regulated by a lncNAT (lncRNAPiRa.31525.1). The lncRNAPiRa.85000.6 lncRNA, which was predicted to target an ethylene receptor 2 (*ETR2*) gene involved in the ethylene signal transduction pathway, was transcribed in the same strand and orientation than its RNA partner from an upstream location. In addition, two genes encoding for photoassimilate-responsive protein 1 (PAR1) were predicted to be targeted by lncRNAPiRa.61651.3 and lncRNAPiRa.33277.3, the latter being DE between conditions.

The pine lncRNA lncRNAPiRa.79902.12 was predicted to target two genes encoding for the pyruvate decarboxylase 1 (PDC1) enzyme, which both were up-regulated by the pathogen infection. Furthermore, one gene that participates in chromatin modifications (chromatin remodelling 24) and three genes that contain canonical RNA-binding domains (pentatricopeptide repeat-containing protein, ribosomal RNA methyltransferase FtsJ domain containing protein, CCCH-type Znf protein) were predicted to be targeted in an antisense manner by lncRNAs. None of the three genes belonged to the DEGs.

The enrichment analysis of GO terms and KEGG pathways of the nearby protein-coding RNAs revealed potential functions in which DE lncRNAs could be involved (**Figure 5**). The three target genes regulating the down-regulated lncRNAs were not associated to any GO term neither KEGG pathway, thus the analysis showed results only for the up-regulated lncRNAs (**Table S12**). Biological and metabolic processes were the most representative GO terms for the biological process category, followed by macromolecule metabolic process and response to stimulus and stress in this dataset. Several GO terms associated with low-oxygen conditions including response to hypoxia and response to decreased oxygen levels were enriched. In addition, catabolism and metabolism of allantoin were also enriched. Genes involved in cell periphery and cell wall were represented for cellular components. For molecular functions, the pine lncRNAs were enriched for GO terms such as catalytic activity, binding and hydrolase activity. The KEGG pathways enriched in the target genes of the up-regulated lncRNAs were ‘glycolysis/gluconeogenesis’ and ‘microbial metabolism in diverse environments’ (**Table S13**).

**Figure 5.**
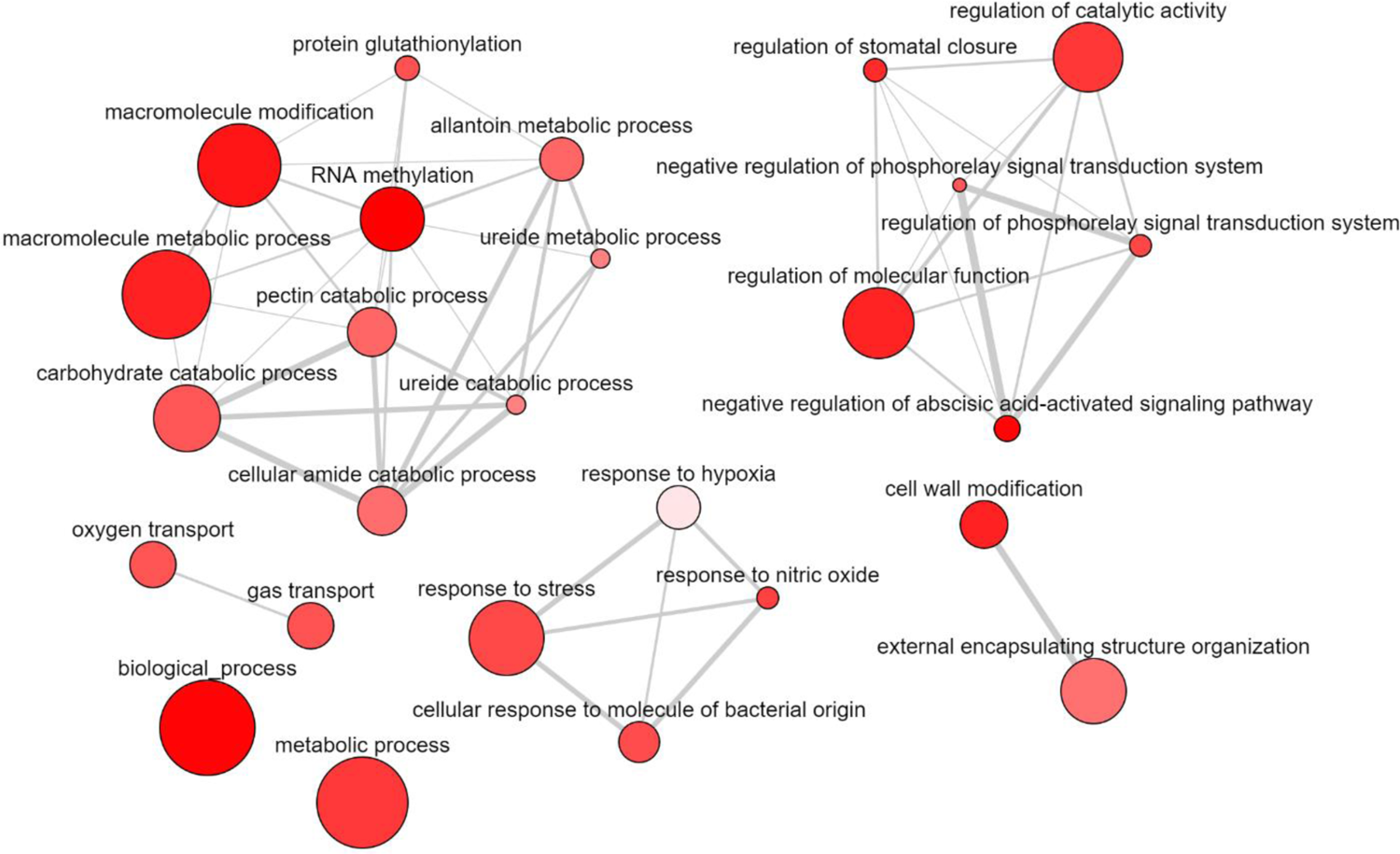
Enriched GO terms visualization of the DE lncRNA targeted genes constructed by REVIGO. Connections are based on the structure of the GO hierarchy. The colour of the bubble reflects the p-value obtained in the functional enrichment analysis, while its size indicates the frequency of the GO term in the underlying UniProt-GO Annotation database. Highly similar GO terms are linked by edges in the graph, where the line width indicates the degree of similarity.

### 3.6. Co-expression gene modules associated with *P. radiata* defence response

A dendrogram, in which the samples were clustered according to their condition using the CEMiTool package, was generated (**Figure 6A**). The modular expression analysis revealed genes that may act together or are similarly regulated during the defence responses to *F. circinatum* infection. The dissimilarity threshold of 0.8 was used as a cut-off on hierarchical clustering, which identified two co-expression modules (**Figure 6B-7C**). The largest module contained 320 co-expressed transcripts (M1): 307 DEGs, 13 DE lncRNAs, and three targeted genes (PDC1, PME and RLK) (**Table S14**).

**Figure 6.**
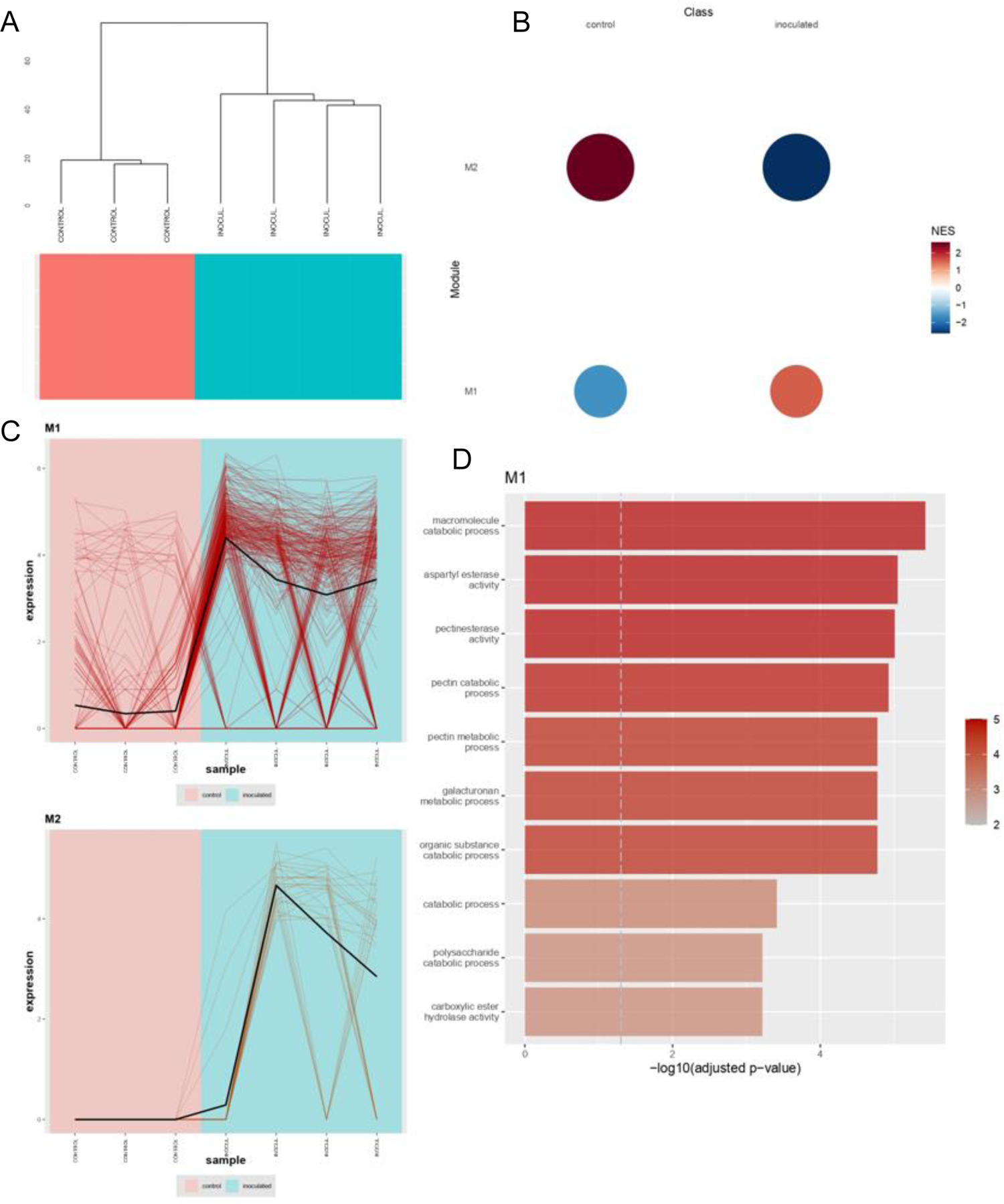
Two co-expression modules were identified among the DE lncRNAs, DEGs and targeted genes using CEMiTool package. (A) Dendrogram of samples clustered according to their condition. (B) Gene set enrichment analysis (GSEA)-based identification of two gene co-expression modules. Red coloring denotes a positive NES score, while blue coloring denotes a negative NES score. (C) Expression profiles for both expression modules (M1, M2). Each line represents a transcript and its change in expression across conditions. (D) Barplot for top GO terms enriched in M1 module. *x*-axis and colour transparency display -log_10_ of the Benjamini-Hochberg (BH)-adjusted p-value. Dashed vertical line indicates BH-adjusted p-value threshold of 0.05.

Transcripts in M1 were enriched mainly for biological processes related to the pectinesterase activity and cell wall remodeling among others (**Figure 6D**) (**Table S15**). Indeed, three DEGs encoding for pectin methylesterase 17 were identified as gene hubs in this module (**Table 3**). The second module (M2) consisted of 30 DEGs and one DE lncRNA (**Table S14**), however, no significant GO terms were identified. The top gene hubs of both modules are shown in **Table 3**.

**Table 3.**
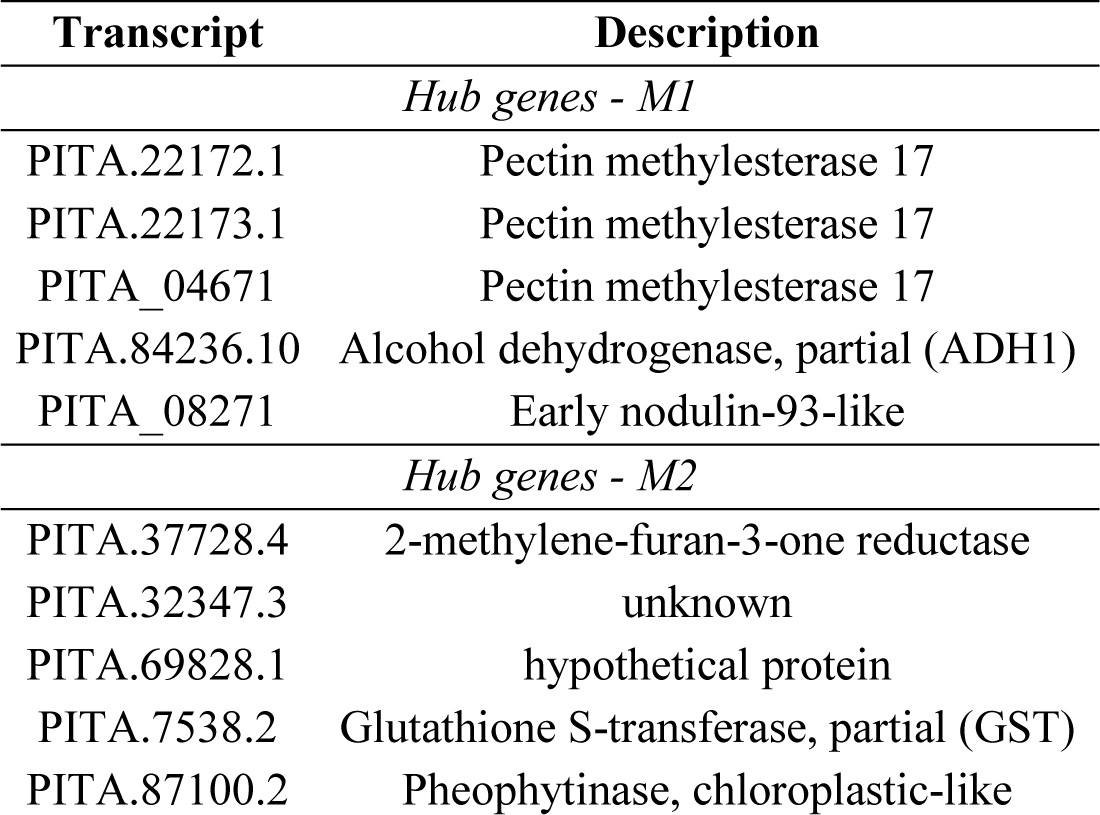
Potential gene hubs of each co-expression gene module.

| Transcript | Description |
| --- | --- |
| *Hub genes - M1* |  |
| PITA.22172.1 | Pectin methylesterase 17 |
| PITA.22173.1 | Pectin methylesterase 17 |
| PITA_04671 | Pectin methylesterase 17 |
| PITA.84236.10 | Alcohol dehydrogenase, partial (ADH1) |
| PITA_08271 | Early nodulin-93-like |
| *Hub genes - M2* |  |
| PITA.37728.4 | 2-methylene-furan-3-one reductase |
| PITA.32347.3 | unknown |
| PITA.69828.1 | hypothetical protein |
| PITA.7538.2 | Glutathione S-transferase, partial (GST) |
| PITA.87100.2 | Pheophytinase, chloroplastic-like |

## 4. Discussion

Over the past decade, the complexity of eukaryote genome expression has become apparent mainly due to the development of next-generation sequencing technologies. Particularly, the sequencing of RNA (RNA-Seq) has revealed an important part of non-coding transcriptome that should not be ignored. Indeed, a large number of studies have recently reported lncRNAs to be essential in the regulation of a wide range of biological and molecular processes by activating their nearly protein-coding genes using a *cis*– mediated mechanism or distant genes in a *trans*-acting manner (Geisler and Coller 2013). Stress conditions lead to transcriptomic reprogramming where lncRNAs also play a key role. In plants, numerous lncRNAs under biotic stress have been identified to date, although further studies for non-model plants are still required. In the last years, the transcriptomic responses of conifers to fungal infections have been increasingly studied. In particular, several transcriptomic studies have demonstrated that the *F. circinatum* infection causes substantial changes in the pine gene expression (Visser *et al*. 2015, 2018, 2019; Carrasco *et al*. 2017; Hernandez-Escribano *et al*. 2020; Zamora-Ballesteros *et al*. 2021). However, to our knowledge, no reports investigating the long non-coding RNAs of conifer trees in response to fungal attacks have been published so far. The results reported here, therefore, provide a first insight into the regulatory mechanisms of lncRNAs involved in defence reactions against *F. circinatum* of a highly susceptible species such as *P. radiata*.

The combination of the strand-specific RNA-Seq approach and high coverage sequencing (up to 84 million reads per sample) allowed the identification of lncRNAs that are commonly expressed at low levels and lncNATs that would otherwise have been difficult to find (Rai *et al*. 2019). Overall, a total of 13,312 lncRNAs were identified from the *P. radiata* transcriptome, of which 164 were *F. circinatum*-responsive lncRNAs comprised mainly by intergenic lncRNAs. This is consistent with previous analyses where the number of lncRNAs in response to a biotic stress was comparable. In *Paulownia tomentosa*, two similar studies found 112 and 110 lncRNAs to be involved in phytoplasma infection (Wang *et al*. 2017; Fan *et al*. 2018). Similarly, among 94 and 302 lncRNAs were identified in susceptible and resistant *M. acuminata* roots in response to *F. oxysporum* f. sp. *cubense*, with the highest value in the resistant roots after 51 hours post-inoculation (Li *et al*. 2017). The number of *S. sclerotiorum*-responsive lncRNAs was slightly higher in *B. napus* with 662 at 24 h decreasing until 308 at 48h (Joshi *et al*. 2016). In addition, intergenic lncRNAs were also the most abundant responsive transcripts in all these studies. Therefore, the pattern appears to follow the same trend in conifer trees.

In general, lncRNAs demonstrate low and tissue-specific expression patterns and lack of conservation (Quan *et al*. 2015; Yu *et al*. 2019; Chen *et al*. 2020). Indeed, lncRNAs of *P. radiata* showed lower expression than the protein-coding RNAs, and only 19.3% of them were conserved among 46 non-conifer plant species. However, the low level of transcriptome conservation in *P. radiata* to angiosperms has also been shown in xylem tissues (15-32%; E-value ≤ 10^-5^), compared with the highly conserved xylem transcriptome within conifers (78-82%; E-value ≤ 10^-5^) (Li *et al*. 2010). Thus, it may not be a characteristic of conifer lncRNAs. The genomic features of the lncRNA transcripts of *P. radiata* were consistent with those previously characterized in other organisms (Cabili *et al*. 2011). As expected, the lncRNAs were shorter in terms of overall length and contained lower number of exons. The length of the exons was also shorter in lncNATs and intronic lncRNAs when comparing with protein-coding RNAs, however, the distribution of the length of exons belonging to lincRNAs was closer to that of the protein-coding transcripts. In this regard, some exceptions have been found in other plants such as cotton (*Gossypium arboretum*) and chickpea (*Cicer arietinum*) where the exon length of the lincRNAs were even longer than protein-coding RNAs (Zaynab *et al*. 2018). The GC content of the assembled transcripts of *P. radiata* (43.1%) was similar to that of the transcriptome of other *Pinus* species such as *P. tecunumanii* (44%) (Visser *et al*. 2018). Separately, the GC content in pine lncRNAs (41%) was lower than in protein-coding RNAs (44.8%), which had been reported before as a common feature of lncRNAs due to different evolutionary pressures in ORFs (Shuai *et al*. 2014).

The role of lncRNAs in the positive or negative regulation of gene expression is well known (Quan *et al*. 2015). One of the conserved mechanism of action of the lncRNAs is their function as decoys by sequestering RNA-binding proteins (RBP), miRNAs or chromatin-modifying complexes (Wang and Chang 2011). Thus, the lncRNA ultimately inhibits its particular function. Several DE lncRNAs of *P. radiata* inoculated by *F. circinatum* seem to fit into this functional mechanism. Four antisense lncRNAs were predicted to target genes encoding RBPs including pentatricopeptide repeat-containing protein (PPR2), ribosomal RNA methyltransferase FtsJ domain containing protein, CCCH-type zinc finger protein and RNA recognition motif (RRM) containing protein. Moreover, another antisense DE lncRNA was predicted to target a chromatin-remodelling gene. Therefore, the reprogramming exerted by the infection of *F. circinatum* on pine transcription affects not only the protein-coding genes, but also the non-coding part of the genome.

The induction of plant defences is a complex biological process that causes a dramatic transcriptomic reprogramming throughout the genome (Kovalchuk *et al*. 2013). Previous studies have shown that a vast number of genes are either up- or down-regulated in response to *F. circinatum* infection (Carrasco *et al*. 2017; Visser *et al*. 2019; Hernandez-Escribano *et al*. 2020; Zamora-Ballesteros *et al*. 2021). Several functional groups of genes have repeatedly been identified as induced upon the pathogen infection. These groups include signal perception and transduction, biosynthesis of defence hormone and secondary metabolites, and cell wall reinforcement and lignification. Some of the GO terms enriched by the lncRNAs identified in this study were related to these functional groups including biological processes such as cell wall modification and signalling of the abscisic acid, ethylene and cytokinin hormones. These results suggest for the first time that the lncRNAs may play a key role in the process of pine defence to *F. circinatum* as previously reported in other pathosystems (Zhu *et al*. 2014; Sanchita *et al*. 2020).

Plant signalling molecules such as protein kinases, reactive oxygen species (ROS) and hormones are critical in mounting an appropriate defence response (Yu *et al*. 2017). Genes with kinase activity have a role in signal transduction triggering the downstream signalling. Two genes with predicted functions in receptor-like kinase were *cis*-regulated by lncRNAs, being one of them DE by the pathogen infection. The other one was potentially regulated by a lncNAT. Positive *cis*-regulatory feature of NATs by mediating histone modifications at the locus has been previously reported (Yu *et al*. 2019). This behaviour has been also seen in LAIR, a rice lncNAT that up-regulates the expression of its neighbour leucine-rich repeat receptor kinase (Wang *et al*. 2018). Despite that a large number of genes (43) encoding glutathione S-transferases (GSTs) were up-regulated under the pathogen infection, the GST predicted to be regulated by the downstream lncRNAPiRa.64704.1 was not among the DEGs. Joshi *et al*. (2016) also identified one lncRNA of *B. napus* located in the upstream of a gene encoding for a GST in response to *S. sclerotiorum* infection. GST genes are highly induced under biotic stress due to their role in the attenuation of oxidative stress and the participation in hormone transport (Gullner *et al*. 2018). In addition, a transcript predicted to encode a non-symbiotic hemoglobin 1, which is involved in ROS and NO scavenging (Bahmani *et al*. 2019), was DE in the analysis and predicted to be targeted by lncRNAPiRa.19024.1. These findings seem to indicate that lncRNAs could be also involved in the cell detoxification after an oxidative burst provoked by a fungal infection.

Phytohormones trigger an effective defence response against biotic stress (Checker *et al*. 2018). Several studies have pointed to lncRNAs as participants in the complex network of hormone regulation. In *M. acuminata* infected by *F. oxysporum* f. sp. *cubense*, lncRNAs were found to be predominantly associated with auxin and salicylic acid signal transduction in susceptible cultivars, whereas all phytohormones were potentially regulated by lncRNAs in resistant cultivars (Li *et al*. 2017). Genes related to the salicylic acid-mediated defence process were co-expressed with lncRNAs in kiwifruit plant challenged with the bacteria *P. syringae* (Wang *et al*. 2017). Likewise, lncRNAs of resistant walnuts to *C. gloeosporioides* were predicted to *trans*-regulate genes involved in defence pathways of the jasmonic acid and auxins (Feng *et al*. 2021). A previous transcriptome analysis of *P. radiata* showed the induction of abscisic acid signalling under the infection of *F. circinatum* (Carrasco *et al*. 2017). A type 2C protein phosphatase (PP2C) family gene, which negatively regulates abscisic acid responses (Cao *et al*. 2016; Jung *et al*. 2020), could be regulated by lncRNAPiRa.47042.1 located upstream in the same strand despite not belonging to the DEGs. The implication of this lncRNA in the abscisic acid signalling regulation would need further investigation.

The phytohormone ethylene represents one of the core components of the plant immune system (Müller and Munné-Bosch 2015). When ethylene binds with its ETRs activates the transcriptional cascade of ethylene-regulated genes (Sakai *et al*. 1998). Seedlings of *P. tecunumanii*, *P. patula*, *P. pinea* and *P. radiata* inoculated with *F. circinatum* have demonstrated to induce ethylene biosynthesis and signalling genes (Carrasco *et al*. 2017; Visser *et al*. 2019; Zamora-Ballesteros *et al*. 2021); however, only *ETR2* has been found to be induced in the moderate resistant specie *P. pinaster* at 5 and 10 dpi (Hernandez-Escribano *et al*. 2020). Under stress conditions, when the concentration of ethylene is high, the transcription of *ETR2* contributes to the stabilization of ethylene levels by attenuating its signalling output and restore the ability to respond to subsequent ethylene signal (Zhao and Guo 2011). In the present study, *ETR2* has not been DE in *P. radiata* but was presumably influenced by lncRNAPiRa.85000.6, which has been DE by *F. circinatum*. Therefore, we can hypothesize that the ethylene response seems to be fine-tuned in *P. pinaster*, which does not occur in *P. radiata*, possibly due to the influence of this lncRNA located upstream of its transcription. It would be worthwhile to further investigate the regulatory function of this lncRNA as it could be a key factor in overcoming the PPC disease.

The potential function of lncRNAs in wood formation has been previously observed in different plant species. In a study of cotton lncRNAs, these were enriched for lignin catabolic processes and their role in lignin biosynthesis by regulating the expression of LAC4 was suggested (Wang *et al*. 2015). In *Populus*, 16 genes targeted by lncRNAs were involved in wood formation processes, including lignin biosynthesis (Chen *et al*. 2015), and 13 targeted genes were associated to cellulose and pectin synthesis (Tian *et al*. 2016). In addition, the lncRNA NERDL regulates the Needed for rdr2-independent DNA methylation (*NERD*) gene, which is also involved in the wood formation in *Populus* (Shi *et al*. 2017). The enzyme that catalyse the hemicellulose xyloglucan was predicted to be targeted by a lncRNA of *Paulownia tomentosa* and had a role in the hyperplasia caused by a phytoplasma infection (Zhe Wang *et al*. 2017). Cell wall reinforcement and lignification are the most common induced defences against pathogens, for that, the cell wall suffers a remodelling process that has been documented in the *P. radiata*-*F. circinatum* pathosystem (Carrasco *et al*. 2017; Zamora-Ballesteros *et al*. 2021). The demethylesterification of pectin, controlled by PMEs, is considered to affect the porosity of the cell wall and, thus, exposes the plant to an easier degradation by pathogen enzymes (Raiola *et al*. 2011). However, PME activity has been also associated with the activation of plant immunity and resistance against pathogens (Del Corpo *et al*. 2020). In a recent study, in contrast to *P. radiata*, the resistant species *P. pinea* infected by *F. circinatum* showed a high induction of pectin methylesterase inhibitor (PMEI) genes and an inhibition of PMEs (Zamora-Ballesteros *et al*. 2021). In the present study, two lncRNAs were predicted to target two PMEs, one of them was up-regulated by the pathogen infection, which could suggest a positive regulation from the lncRNA activity. In addition, the co-expression analysis of *F. circinatum* responsive lncRNAs and mRNAs indicated a clear enrichment for PME activity. The transcriptional regulation of these enzymes could be related to the susceptibility of *P. radiata* and would be worth further investigation. Another gene containing a cellulase domain was also up-regulated in the expression analysis of protein-coding RNAs and predicted to be regulated by an induced lncRNA. Moreover, the analysis identified a potential lncRNA *cis*-regulating positively a gene encoding for 4CL3, one of the key enzymes of the phenylpropanoid pathway. In plants, this pathway leads to the production of secondary metabolites and cell wall lignification, both associated to plant defence. The transcriptional regulation of the *4CL* gene by lncRNAs has been also reported in *P. tomentosa*, that together with the targeted gene encoding the caffeoyl-CoA 3-O-methyltransferase (CCOMT) enzyme by another lncRNA, highlighted the potential role of these molecules in lignin formation in wood with different properties (Chen *et al*. 2015). These findings provide increasing evidence for the involvement of lncRNAs in cell wall remodelling and lignification process.

Although the role of the hypoxia in the plant-pathogen interaction has not yet been determined, hypoxia-responsive genes have been reported to be induced in some plants during pathogen infections (Loreti and Perata 2020). Indeed, the analysis of DEGs showed that a large number of genes encoding for PDC1 and alcohol dehydrogenase 1 (ADH1), which are required in the fermentative pathway under low-oxygen conditions, were highly induced by *F. circinatum* infection (>10 log_2_[fold change]; **Table S8**). Among them, two PDC1 were potentially targeted by two pine lncRNAs. This together with the functional analysis results of the lncRNAs where several enriched GO terms were associated to hypoxia suggests a role of pine lncRNAs in an insufficient oxygen situation.

In summary, the computational analysis allowed to identify 13,312 lncRNAs in *P. radiata*. Compared to the protein-coding RNAs, the lncRNAs were shorter, with fewer exons and showed lower expression levels. In total 164 lncRNAs were reported as responsive to *F. circinatum* infection. GO enrichment of genes that either overlap with or are neighbours of these pathogen-responsive lncRNAs suggested involvement of important defence processes including signal transduction and cell wall reinforcement. These results present a comprehensive map of lncRNAs in *P. radiata* under *F. circinatum* infection and provide a starting point to understand their regulatory mechanisms and functions in conifer defence. In turn, a thorough understanding of the mechanism of gene regulation will contribute to the improvement of breeding programs for resistant pine commercialization, one of the most promising approaches for PPC management.

