## Supplementary figures for "Genome-Wide Identification and Characterization of *Fusarium circinatum*-Responsive lncRNAs in *Pinus radiata*"

|  | Rate |
| --- | --- |
| Chenopodium quinoa | 1.24957% |
| Zea mays | 0.715547% |
| Vitis vinifera | 2.37781% |
| Populus trichocarpa | 5.03101% |
| Prunus persica | 1.36511% |
| Malus domestica | 2.05969% |
| Ananas comosus | 1.48981% |
| Brassica napus | 1.39883% |
| Arabidiopsis thaliana | 1.28059% |
| Oryza rufipogon | 0.477536% |
| Amborella trichopoda | 1.8954% |
| Glycine max | 1.55479% |
| Arabidopsis lyrata | 1.146% |
| Brachypodium distachyon | 0.179083% |
| Capsella grandiflora | 0.573888% |
| Capsella rubella | 0.685871% |
| Carica papaya | 0.401304% |
| Citrus sinensis | 1.60031% |
| Citrus clementina | 1.41423% |
| Cucumis sativus | 1.86625% |
| Eucalyptus grandis | 0.970037% |
| Eutrema salsugineum | 0.419413% |
| Fragaria vesca | 0.314017% |
| Gossypium raimondii | 1.25712% |
| Linum usitatissimum | 1.57814% |
| Manihot esculenta | 1.12994% |
| Medicago truncatula | 0.651095% |
| Mimulus guttatus | 1.75739% |
| Musa acuminata | 3.41439% |
| Ostreococcus lucimarinus | 0.199601% |
| Phaseolus vulgaris | 2.26105% |
| Physcomitrella patens | 0.361197% |
| Ricinus communis | 0.524059% |
| Selaginella moellendorffii | 0.551876% |
| Setaria italica | 0.286369% |
| Solanum lycopersicum | 0.930233% |
| Solanum tuberosum | 1.4521% |
| Sorghum bicolor | 0.301602% |
| Spirodela polyrhiza | 0.900047% |
| Theobroma cacao | 0.609185% |
| Triticum aestivum | 0.592478% |
| Zostera marina | 1.7229% |

**Figure S1.** Homology ratio of *Pinus radiata* lncRNAs to lncRNAs of plant species from the CANTATA and GreeNc databases. The bars represent the number of hits of pine lncRNAs with those of each plant species to the total number of lncRNAs of each plant species.

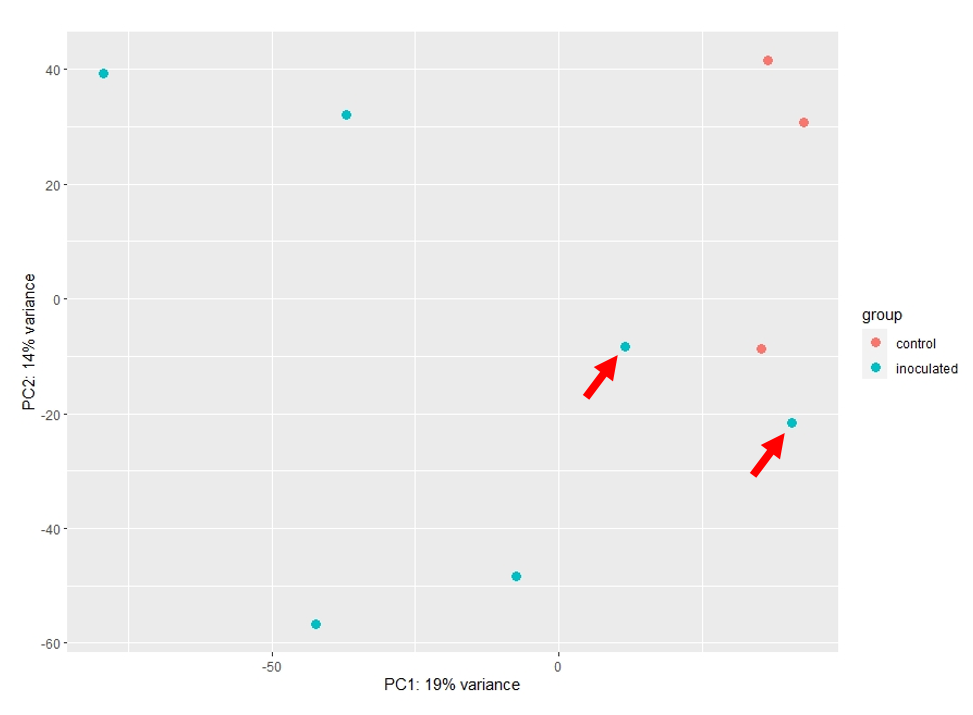

**Figure S2.** Two-dimensional scatterplot of the principal component analyses (PCA) for lncRNA expression data of *Pinus radiata*. Red arrows indicate the discarded samples for the downstream analysis.
